# T Lymphocyte-Specific Deletion of SHP1 and SHP2 Promotes Activation-Induced Cell Death of CD4+ T Cells and Impairs Antitumor Response

**DOI:** 10.1101/2025.01.03.631277

**Authors:** Connor J.R. Foster, Jasper Du, Oscar Pundel, Mitchell J. Geer, Ryan C. Ripert, Jia Liu, Taylor A. Heim, Kiyomi Y. Araki, Amanda W. Lund, Jun Wang, Benjamin G. Neel

## Abstract

SHP1 (PTPN6) and SHP2 (PTPN11) are closely related protein-tyrosine phosphatases (PTPs), which are autoinhibited until their SH2 domains bind paired tyrosine-phosphorylated immunoreceptor tyrosine-based inhibitory/switch motifs (ITIMs/ITSMs). These PTPs bind overlapping sets of ITIM/ITSM-bearing proteins, suggesting that they might have some redundant functions. By studying T cell-specific single and double knockout mice, we found that SHP1 and SHP2 redundantly restrain naïve T cell differentiation to effector and central memory phenotypes, with SHP1 playing the dominant role. Surprisingly, loss of SHP2 alone in T cells enhanced the antitumor effects of anti-PD-1 antibodies, whereas there was no effect of SHP1 deletion. Also unexpectedly, the absence of both PTPs resulted in poorer tumor control and failure to respond to PD-1 blockade, associated with reduced frequency and activation of T cells and dendritic cells. Mechanistic studies revealed that CD4+, but not CD8+ T cells lacking SHP1 and SHP2 show increased activation-induced cell death upon anti-CD3/CD28 stimulation. Adoptive transfer of antigen-specific CD4+ T cells restored normal levels of tumor control in mice lacking both PTPs. Together, our results demonstrate that SHP1 or SHP2 is required to prevent activation-induced cell death of CD4+ T cells and is critical for tumor immunity, raising the possibility that inhibition of SHP2 might augment the therapeutic efficacy of PD-1-based immune therapy.

**SIGNIFICANCE STATEMENT:** SHP1 and SHP2 are related protein tyrosine phosphatases that associate with several of the same ITIM/ITSM-containing receptors or T cell receptor (TCR) signaling molecules. The individual roles of SHP1 and SHP2 in T cells have been reported previously, but potentially redundant functions are less well understood. Here we uncover an essential function in CD4+ T cells that is manifest only in the absence of both enzymes and is critical for the control of tumors.

## INTRODUCTION

The protein tyrosine phosphatases SHP1 (encoded by *PTPN6*) and SHP2 (encoded by *PTPN11*) have high structural similarity (∼53% identify) comprising two SH2 domains at the N-terminus, a PTP domain, and a C-terminal tail (1). SHP1 has an additional sequence in its C-terminus that promotes recruitment to lipid rafts and is important for its inhibitory activity in T cells (2). Under basal conditions, SHP1 and SHP2 are autoinhibited (closed conformation) owing to binding of the non-tyrosine-binding portion (“backside) of the N-SH2 domain to the PTP domain (3, 4). Both PTPs “open” and become active when their SH2 domains bind to tandem immunoreceptor tyrosine-based inhibitory motifs (ITIMs; consensus sequence S/I/V/LxYxxI/V/L) or immunoreceptor tyrosine-based switch motifs (ITSMs; consensus sequence TxYxxV/I) (1, 5). SHP1 and SHP2 bind a variety of proteins, including receptor tyrosine kinases (RTKs), cytokine receptors, scaffolding adaptors, and inhibitory receptors. Some of these proteins bind only one of the SHPs, while others bind both, raising the possibility of that the SHPs might have some redundant functions.

Several early studies used *motheaten* or *motheaten viable* mice to characterize the role of SHP1 in T lymphocytes. However, due to the effects of SHP1 deficiency on other lymphohematopoietic cells, it is unclear which of these effects seen in these mice are T cell-autonomous. Our group and others have used conditional deletion to study SHP1 in thymocytes and peripheral T cells. SHP1 inhibits TCR signaling during thymocyte development; consequently, *Ptpn6* deletion leads to fewer thymocytes surviving negative selection (6). SHP1 also inhibits TCR signaling in peripheral T cells, lowering the threshold for proliferation and limiting total expansion (7–9). Owing to its role as a negative regulator of IL-4 signaling, SHP1 restricts differentiation of naïve T cells to effector and central memory phenotypes and limits Th2 polarization of CD4+ T cells (10).

SHP2 is expressed ubiquitously, and global deletion in mice is embryonic lethal (11, 12). SHP2 has a well-documented positive role in RAS-RAF-MEK-ERK signaling, acting upstream of SOS1/2 (13). Two previous reports indicate that SHP2 has no impact on thymocyte development (14, 15). However, SHP2 reportedly limits expansion of and cytokine secretion by mature antigen-specific CD8+ T cells *in vivo* (14). SHP2 is also reported to associate with the CD3ζ chains and SLP76 upon stimulation in Jurkat T cells (16, 17). However, it appears to be individually dispensable for T cell intracellular Ca^2+^ release and CD3ζ chain phosphorylation (18). SHP2 also binds directly to some cytokine receptors and to the scaffolding adaptors GAB1/2 and therefore could impact multiple pathways relevant in T cells (13, 19).

PD-1 is a critical inhibitory receptor that limits antitumor immune responses, and anti-PD-1 antibodies have had major therapeutic impact in multiple malignancies. Early studies suggested a role for SHP2 in PD-1 signaling (20, 21). In primary human CD4+ T cells, PD-1 binds SHP1 and SHP2 (21), although quantitative mass spectrometry of PD-1 immunoprecipitates from primary murine T cells revealed preferential binding to the latter SHP (22). PD-1-bound SHP2 dephosphorylates CD28 and, to a lesser extent, LCK *in vitro* (23). However, functional studies interrogating the role of SHP2 in PD-1 signaling are, at best, conflicting. For example, co-cultures of PD-L1-overexpressing Raji cells and PD-1-overexpressing Jurkat cells have yielded mixed results. One study identified both SHP1 and SHP2 as mediators of PD-1 action, while another concluded that SHP2 is the primary mediator, with a minor contribution from an unidentified signaling component (22, 24). A prior report found that mice with a T cell-specific deletion of *Ptpn11* responded to systemic PD-1 blockade in an MC38 tumor model, apparently suggesting that SHP2 was not required for PD-1 signaling in T cells (14). However, more recent work argues that in mice, PD-1 action in the myeloid cell compartment is more important for its anti-tumor effect against MC38 tumors (25, 26). While all published *in vitro* studies suggest that SHP2 mediates PD-1 signaling, it remains unclear whether SHP1 does so as well (22–24). Besides PD-1, multiple inhibitory receptors expressed on T cells bind SHP1 and SHP2 to varying degrees, including BTLA, CEACAM1, LAIR1, and, in humans, LILRB1 (27–29). Consequently, the SHPs might redundantly restrict T cell activation downstream of these molecules. Here, we investigate the effects of combined *Ptpn6/Ptpn11* deletion on T cell development and anti-tumor response in the presence and absence of PD-1 blockade.

## RESULTS

### Effect of SHP1 and SHP2 on Thymocyte Development

To dissect the roles of SHP1 and SHP2 (SHPs) in T cell biology, we generated mice that were hemizygous for *Cd4-Cre* and homozygous for floxed *Ptpn6* (SHP1 knockout/KO) and/or *Ptpn11* (SHP2 knockout/KO) alleles. We validated that splenic T cells from *Cd4-Cre Ptpn6^fl/fl^ Ptpn11^fl/fl^* (double knockout/DKO) mice lacked expression of both PTPs (Fig. S1A). CD4-Cre expression commences at the double positive (DP) stage of thymocyte development (30), so we assessed the effects of SHP1 and/or SHP2 deficiency in DP cells and their progeny, single positive (SP) cells. DKO mice had fewer total thymic cells compared with their *CD4-Cre Ptpn6^+/+^ Ptpn11^+/+^* (control) counterparts, mainly due to SHP1 deficiency (Fig. S1B). Positive and negative selection occur during the DP stage, and T cell receptor (TCR) signal strength is a critical determinant of survival during selection. Signal strength that is too weak or too strong results in thymocyte death (31). Notably, SHP1 KO mice had significantly fewer DP thymocytes than controls (Fig. S1C), possibly resulting from increased DP thymocyte deletion and consistent with a report that SHP1 deficiency results in fewer DP thymocytes surviving negative selection (6). Deficiency of SHP2 alone had no effect on total number of DP thymocytes, but deficiency of both SHPs resulted in a further decrease compared with SHP1 deficiency.

CD5 and CD69 expression on DP thymocytes correlate with TCR signal strength (32, 33). Consistent with previous reports of increased TCR signal strength in SHP1-deficient thymocytes, our SHP1 KO mice showed increased percentages of CD5^hi^ and CD69+ DP thymocytes compared with control mice (Fig. S1D and E) (6). DKO mice also had an increased CD5^hi^ fraction, although they did not have an increased CD69+ fraction of DP thymocytes compared with control mice. CD4+ and CD8+ single positive (SP) thymocyte numbers were decreased in both SHP1 KO and DKO mice compared with controls (Fig. S1F and G), consistent with fewer thymocytes surviving selection. These data collectively suggest a modest role of SHP1/SHP2 in thymic T cell development associating with TCR signal strength, with a more dominant function from SHP-1.

### SHP1 and SHP2 Redundantly Regulate T Cell Phenotypic Differentiation

While control and SHP2 KO mice had similar splenic T cell numbers, DKO mice had significantly fewer (Fig. 1A). There was also a trend toward lower numbers of splenic T cells in SHP1 KO mice; in the absence of an outlier, SHP1 KO mice also would have significantly fewer T cells than controls. Frequencies of CD4+ T cells in SHP1 KO, SHP2 KO, and DKO mice did not significantly differ from that of controls (Fig. 1B). We showed previously that SHP1 restrains differentiation of naïve T cells to effector/central memory cells via negative regulation of the IL-4 receptor (10). As expected, SHP1 KO mice had fewer naïve (CD62L^hi^ CD44^lo^) splenic CD4+ T cells with a concomitant increase in effector (CD62L^lo^ CD44^hi^) CD4+ T cells (Fig. 1C-E). SHP2 KO alone did not affect naïve, effector, and central memory (CD62L^hi^ CD44^hi^) CD4+ populations. However, combined deficiency of both SHPs accentuated the effects of SHP1 deficiency: DKO mice had significantly fewer naïve CD4+ T cells than SHP1 KO mice and significantly increased splenic central memory CD4+ T cells. SHP1 deficiency led to decreased frequencies of CD8+ T cells, and DKO mice had similarly decreased CD8+ T cell proportions (Fig. 1F). SHP1 KO mice had significantly fewer naïve CD8+ T cells relative to control mice with increases in both the effector and central memory populations (Fig. 1G-I). Again, SHP2 deficiency had no independent effect on differentiation, but combined SHP deficiency accentuated the effects of SHP1 deficiency with DKO mice having even more effector and central memory CD8+ T cells.

**Figure 1.**
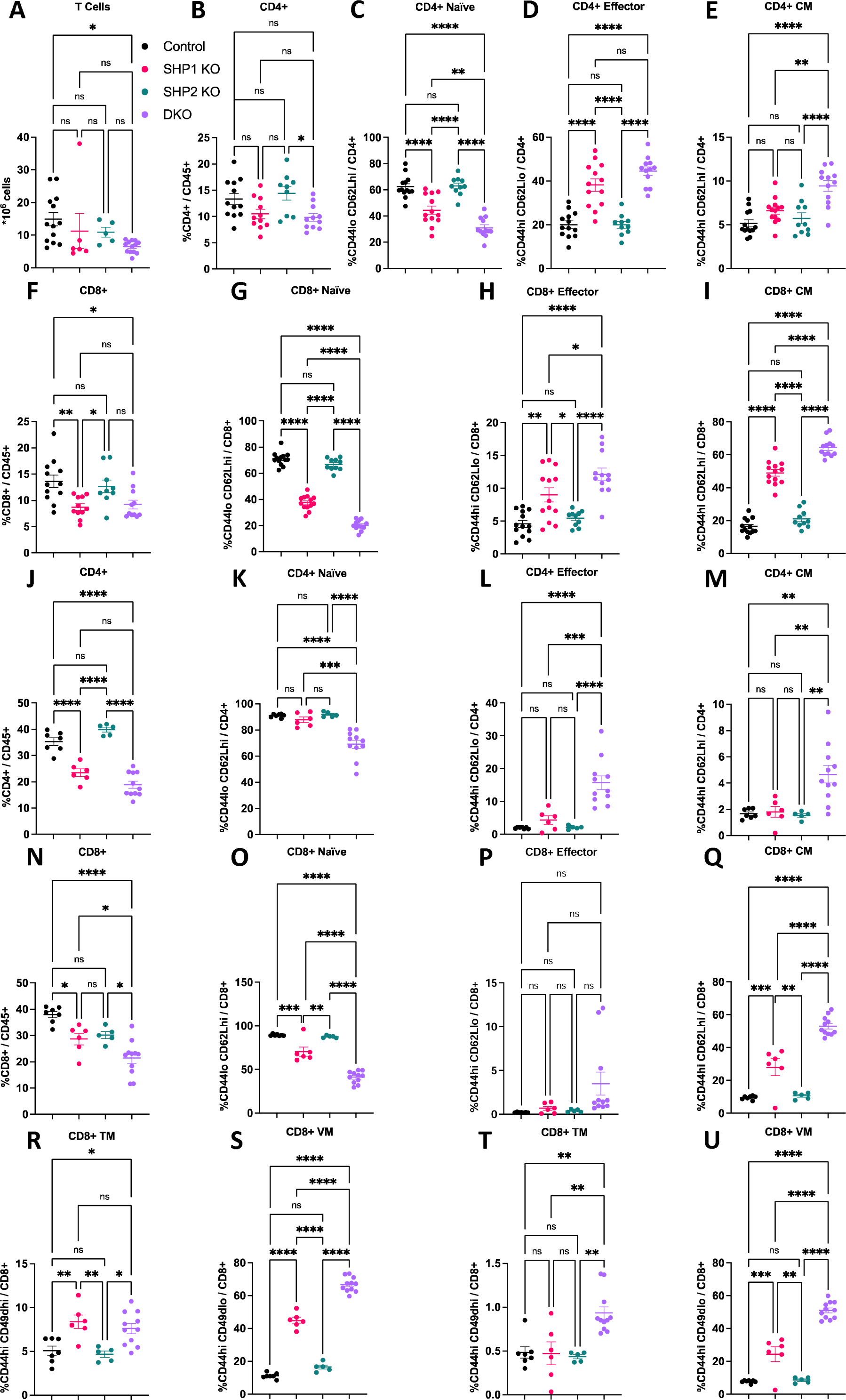
SHP1 and SHP2 redundantly restrict naïve T cell differentiation. (A-I) Splenic T cell populations from the indicated mice, CM; central memory. (J-Q) Inguinal lymph node T cell populations. (R-S) Splenic memory T cell subpopulations. (R) CD8+ true memory (TM; CD44^hi^CD49d^hi^), (S) CD8+ virtual memory (VM; CD44^hi^CD49d^lo^). (T-U) Inguinal lymph node memory T cell subpopulations. (T) CD8+ TM, (S) CD8+ VM. Data are combined from at least two experiments. Each point represents value from one mouse. Means ± standard error are indicated. Significance was evaluated by one-way ANOVA with multiple comparisons, not significant (ns), *p<0.05, **p<0.01, ***p<0.001, ****p<0.0001

T cell phenotypes in the inguinal lymph nodes (LNs) were similar to those in the spleen. In LNs, CD4+ T cell frequency was significantly reduced in both SHP1 KO and DKO mice compared with control mice, and while the frequency in DKO mice trended lower than in SHP1 KO mice, the difference was not statistically significant (Fig. 1J). Also, unlike in the spleen, LNs from DKO mice had increased percentages of effector and central memory CD4+ T cells relative to control and SHP1 KO mice (Fig. 1K-M). DKO mice had fewer CD8+ T cells in the LNs than SHP1 KO mice, which in turn, had fewer than control mice (Fig. 1N). LN CD8+ T cells of SHP1 KO and DKO mice were skewed toward central memory phenotypes (Fig.1 O-Q). Collectively, these results indicate that SHP1 and SHP2 redundantly restrict T cell differentiation with SHP1 playing the dominant role. The reason why SHP1 KO and DKO mice had fewer T cells in spleen and lymph nodes is unclear but could relate to decreased output from the thymus.

Within the memory compartment, CD44^hi^ CD49d^hi^ CD8+ T cells are reported to be antigen-experienced (true memory; TM), whereas CD44^hi^ CD49d^lo^ CD8+ T cells are antigen-inexperienced (virtual memory; VM) (34). SHP1 KO and DKO mice had small increases in the splenic CD8+ TM compartment, but populations of these cells in all genotypes, including controls, were low (Fig. 1R). Nevertheless, SHP1 deficiency significantly elevated the percentage of CD8+ VM T cells, which increased even further in mice with combined SHP deficiency (Fig. 1S). The TM and VM phenotypes in the LN were consistent with those of the spleen (Fig. 1T, U). Together, these data suggest that, at least in CD8+ T cells, SHP1/SHP2 may also redundantly control the percentage of cells with a memory phenotype that is independent of antigen exposure.

### Either SHP1 or SHP2 Is Required for Antitumor T Cell Responses

PD-1, along with other T cell inhibitory receptors, binds both SHP1 and SHP2. Therefore, we hypothesized that DKO mice would exhibit an enhanced antitumor T cell response against PD-1-responsive tumors compared with control, SHP1 KO, or SHP2 KO mice. We expected that deletion of essential PD-1 signaling mediators would phenocopy PD-1 blockade or deletion, resulting in smaller tumors, regardless of PD-1 antibody treatment. To this end, we injected control, SHP1 KO, SHP2 KO, and DKO mice subcutaneously with MC38 cancer cells and treated them with anti-PD-1 or isotype control antibody. As expected, tumors grew more slowly in control mice treated with anti-PD-1 than in control animals (Fig. 2A and B). Moreover, these mice were more likely to be responders (6/23 responders; see Materials and Methods) than isotype-treated mice (0/20 responders; Fig. 2C). We also found that SHP1 KO mice treated with anti-PD-1 had reduced tumor growth and were more likely to be responders (3/9 responders) compared with isotype-treated mice (0/11 responders; Fig. 2D-F). In contrast to a previous report, we found that SHP1 KO mice were equally (i.e., not less) likely to respond to treatment than control mice (Fig. 2G). SHP2 KO mice treated with anti-PD-1 also had reduced tumor volume over time and were more likely to be responders (16/29 responders) than those treated with isotype antibody (2/23 responders; Fig. 2H-J). To our surprise, PD-1 blockade was actually *more* effective in SHP2 KO mice than in control mice (Fig. 2K). Although only a minority of control mice responded to anti-PD-1 treatment (26%), the majority of SHP2 KO mice were responders (55%). These data indicate that SHP2 restricts T cell-mediated antitumor responses in a T cell-autonomous fashion via a pathway independent of PD-1.

**Figure 2.**
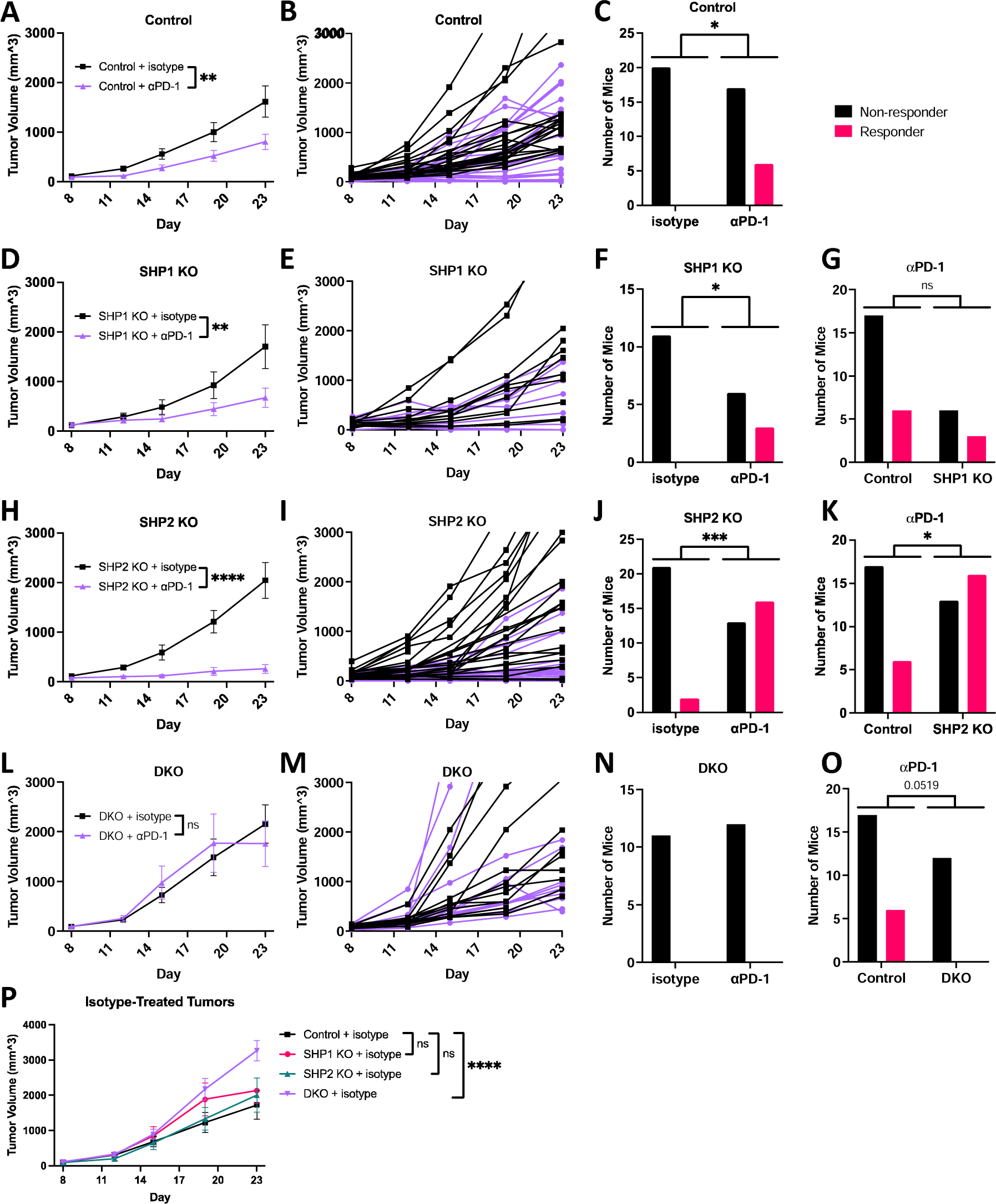
Either SHP1 or SHP2 is required for antitumor T cell responses. (A-P) MC38 tumor growth in mice with isotype (black line) or anti-PD-1 (⍺PD-1; purple line) antibody treatment. (A-C) WT +isotype (n=24), +⍺PD-1 (n=25), (D-F) SHP1 KO +isotype (n=11), +⍺PD-1 (n=10_, (H-J) SHP2 KO +isotype (n=26), +⍺PD-1 (n=29), (L-N) DKO +isotype (n=13), +⍺PD-1 (n=13). (A, D, H, L) Tumor growth, analyzed by two-way ANOVA, (B, E, I, M) Individual tumor growth curves, (C, F, J, N) Histograms of responders and non-responders in isotype-treated versus anti-PD-1-treated mice, analyzed by chi-squared test [except for (N) due to 0 responders in both groups], (G, K, O) Histogram of responders and non-responders in anti-PD-1-treated mice comparing (G) WT and SHP1 KO, (K) WT and SHP2 KO, (O) WT and DKO, analyzed by chi-squared test. (P) Tumor growth in different set of isotype-treated mice analyzed by two-way ANOVA, WT (n=12), SHP1 KO (n=14), SHP2 KO (n=15), DKO (n=13). All comparisons were combined from 2-5 experiments. Mice humanely euthanized before day 23 were excluded from statistical comparisons. Means ± standard error are indicated. Not significant (ns), *p<0.05, **p<0.01, ***p<0.001, ****p<0.0001.

Remarkably, and in contrast to our expectation, isotype- and anti-PD-1-treated DKO mice had similar tumor growth rates, and no DKO mice qualified as anti-PD-1 responders (Fig. 2L-O). Surprisingly, DKO mouse tumors grew significantly larger and faster than tumors from control, SHP1 KO, and SHP2 KO mice, which all had similar tumor growth (Fig. 2P). While our experiments were in progress, Ventura *et al.* also reported that DKO mouse tumors were larger than control tumors (35). These findings collectively suggest DKO T cells are dysfunctional, leading to an impaired antitumor T cell response. Thus, either SHP1 or SHP2 is required for T cell-mediated control of tumor growth and response to PD-1 checkpoint blockade. However, because of the incompetent antitumor response in DKO mice, we cannot determine from the data whether both SHP1 and SHP2 can mediate PD-1 signaling in T cells.

### Combined SHP deficiency in T Cells Leads to a Dysregulated Tumor Immune Microenvironment

We next examined the immune microenvironment in tumors from DKO and control mice at 24 days after injection of MC38 cells. Consistent with growth measurements in Fig. 2, tumors from DKO mice had greater mass than those from control mice (Fig. 3A). Single-cell suspensions from tumors were analyzed by flow cytometry. We found that the proportions of total leukocytes in tumors were similar in both genotypes (Fig. 3B). However, the frequency of CD4+ T cells in DKO mouse tumors was nearly half that of controls (Fig. 3C). CD4+ T cells in tumors from DKO mice and controls showed similar levels of phenotypic exhaustion, as measured by PD-1/TIM3 double positivity (Fig. 3D) or by expression of LAG3 or CTLA4 (Fig. S3A, B). Consistent with reduced IL-4 signaling, we observed polarization of DKO CD4+ T cells away from Th1 and toward Th2 (Fig. S3C-E).

**Figure 3.**
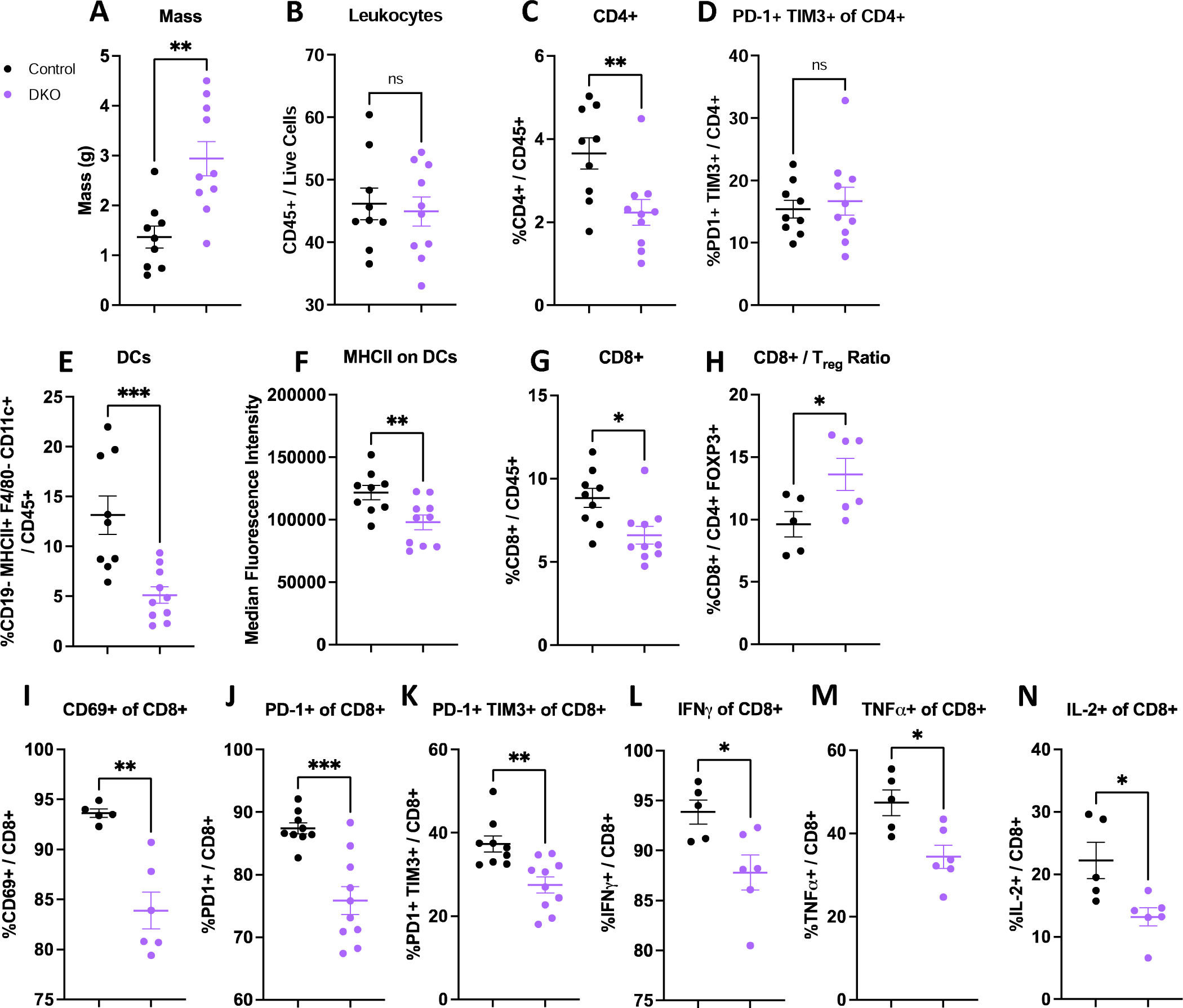
DKO mouse tumors have a dysregulated immune microenvironment. (A-N) Flow cytometric analysis of cells from isotype-treated WT and DKO mouse tumors. (A) tumor mass, (B) %leukocytes, (C) CD4+ T cells, (D) PD1/TIM3 double positive CD4+ T cells, (E) dendritic cells (DCs), (F) MFI of MHC class II on DCs, (G) CD8+ T cells, (H) CD8+ T cell to T_reg_ ratio, (I) CD69+ CD8+ T cells, (J) PD1+ CD8+ T cells, (K) PD1/TIM3 double positive CD8+ T cells. (L-N) Cells suspensions were treated with PMA+ionomycin+Brefeldin A (L) IFNγ+ CD8+ T cells, (M) TNFα+ CD8+ T cells, (N) IL-2+ CD8+ T cells. Data were combined from two experiments. Each point represents value from one mouse. Means ± standard error are indicated. Significance was evaluated by two-tailed unpaired t-tests, not significant (ns), *p<0.05, **p<0.01, ***p<0.001, ****p<0.0001.

We also found that tumors from DKO mice had significantly fewer dendritic cells than those from control mice (Fig. 3E). The intratumoral dendritic cells of DKO mice had significantly lower MHC class II expression than control mice, indicating deficient activation (Fig. 3F). CD8+ T cell numbers were also decreased in DKO mouse tumors (Fig. 3G). The proportion of T_regs_ in the CD4+ T population was similar in control and DKO mouse tumors (Fig. S3F), while the ratio of CD8+ T cells to T_regs_ was actually greater in the latter (Fig. 3H). These results indicate that T_regs_ are unlikely to account for the impaired antitumor responses in DKO mice. The percentages of intratumoral CD8+ T cells expressing CD69 or PD-1 were also significantly decreased (Fig. 3I, J), indicating decreased CD8+ T cell activation. However, the percentage of phenotypically exhausted (PD-1/TIM3 double positive) CD8+ T cells was lower in the tumors from DKO mice (Fig. 3K), arguing against exhaustion as the explanation for their dysfunctional anti-tumor response. CD8+ T cells taken from DKO mouse tumors and stimulated *ex vivo* also had significantly lower production of IFNγ, TNFα, and IL-2 compared with control CD8+ T cells (Fig. 3L-N), highlighting their decreased effector capacity. While DKO mouse tumors had fewer T cells than control mouse tumors, they also had significantly increased proportions of tumor-associated macrophages (TAMs), which had decreased proportions of PD-L1 expression (Fig. S3H and I). These data highlight major differences in the effects of combined SHP1/SHP2 deficiency on CD4+ versus CD8+ T cells, and the consequences of SHP1/SHP2 loss in T cells on tumor-infiltrating dendritic cells and TAMs (see Discussion).

### DKO T Cells Maintain Cytotoxic Capacity

A possible explanation for the impaired DKO antitumor T cell response is that SHP1 or SHP2 is required for T cell cytotoxicity. To generate target cells for testing cytotoxic capacity, we transduced MC38 tumor cells with a construct encoding mouse anti-CD3 Fab fused to the GPI-anchored protein, human CD14 (Fig. S4A). This approach yielded MC38 cells with an outwardly facing, membrane-bound anti-CD3 (MC38-αCD3) to activate murine T cells *ex vivo* in an antigen-independent manner. To test whether DKO T cells could kill target cells, we isolated splenic T cells from control or DKO mice and stimulated them with plate-bound anti-CD3 and anti-CD28 for 48 hours in the presence of 20 ng/mL recombinant IL-2 (Fig. S4B). We cultured these cells for 7 days in IL-2 (in the absence of TCR stimulation), then added them to MC38 or MC38-αCD3 cells and monitored killing. Control and DKO T cells both killed MC38-αCD3 at multiple effector to target (E:T) ratios with more efficient killing of MC38-αCD3 cells than MC38 cells (Fig. S4C, D). Hence, neither SHP1 nor SHP2 is essential for T cell cytotoxicity.

### SHP1 or SHP2 Is Required to Prevent Activation-Induced Cell Death in CD4+ T Cells

Another potential explanation for the dysfunctional antitumor T cell response in DKO mice is defective T cell expansion upon TCR stimulation. To explore this possibility, we stimulated control, SHP1 KO, SHP2 KO, and DKO T cells *ex vivo* with anti-CD3 and anti-CD28 and measured T cell numbers and fractions of CD4+ and CD8+ cells over several days. Control CD8+ T cells expanded more rapidly than CD4+ T cells (Fig. 4A-D), consistent with existing literature (36). CD8+ T cells from all genotypes showed similar net expansion at day 5 post-stimulation. By contrast, while control, SHP1 KO, and SHP2 KO CD4+ T cells expanded similarly over the course of 5 days, DKO CD4+ T cell numbers dropped to near zero (Fig. 4C and D).

**Figure 4.**
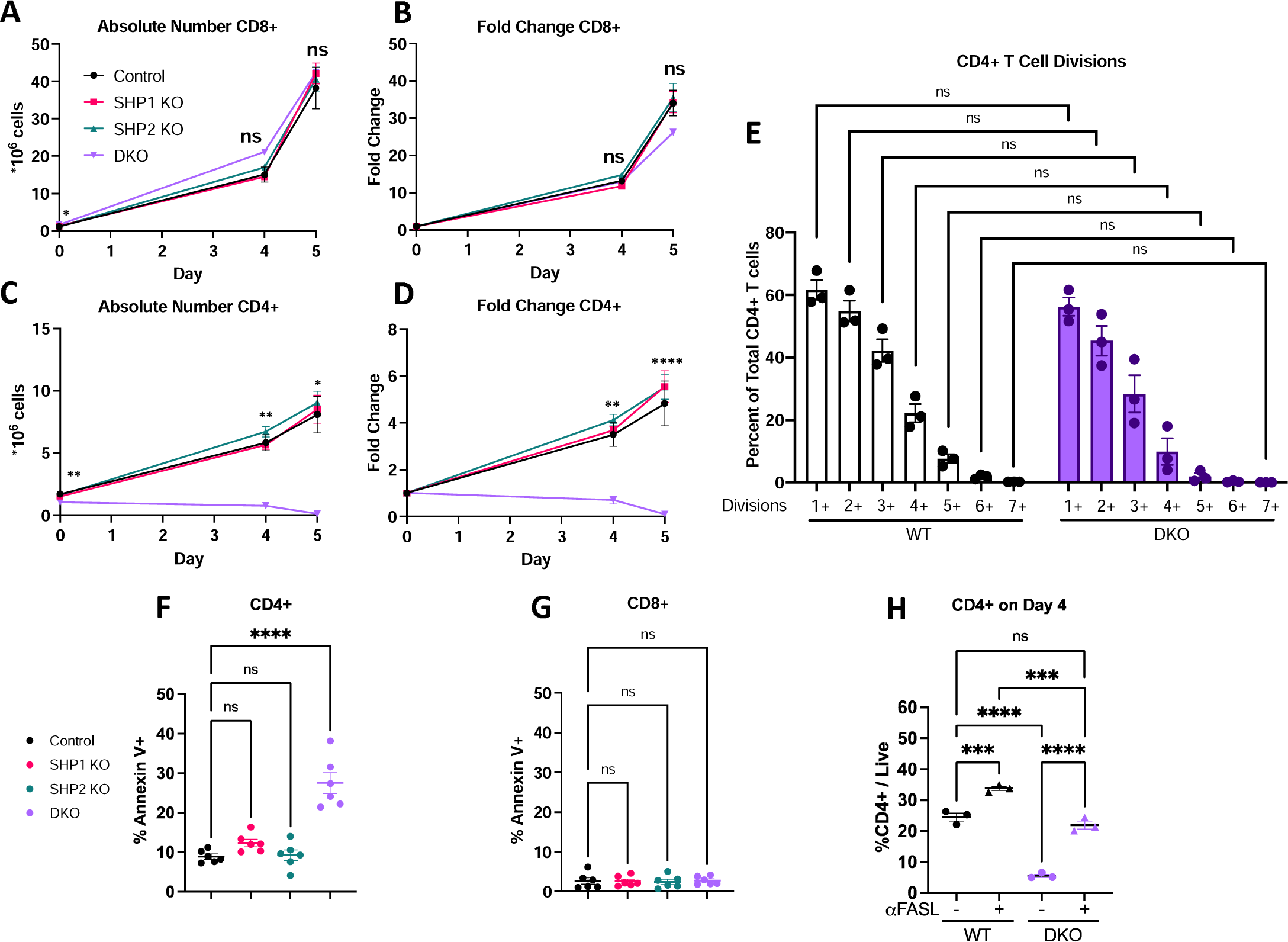
SHP1 or SHP2 is required to prevent activation induced cell death (AICD) in CD4+ T cells. (A-D) T cell proliferation following stimulation with ⍺CD3 and ⍺CD28, (A) Absolute number of CD8+ T cells, (B) Fold change in CD8+ T cell number from baseline, (C) Absolute number of CD4+ T cells, (D) Fold change in CD4+ T cells from baseline. (E) CTV dye dilution assays showing CD4+ T cell divisions after 4 days after stimulation, (F-G) Annexin V staining at 3 hours post-stimulation of (F) CD4+ T cells and (G) CD8+ T cells. (H) Percentage of CD4+ T cells at day 4 post-stimulation with/without FASL blockade. Data are representative of or combined from at least two experiments. Each point represents value from one mouse. Means ± standard error are indicated. Significance was evaluated by one-way ANOVA with multiple comparisons, not significant (ns), *p<0.05, **p<0.01, ***p<0.001, ****p<0.0001.

Defective CD4+ T cell expansion could result from impaired cell proliferation and/or increased cell death. To assess the proliferation of DKO CD4+ T cells, we stimulated CTV-stained T cells and measured dye dilution. At day 4 post-stimulation, similar fractions of control and DKO CD4+ cells had undergone between 1 and 7+ divisions (Fig. 4E), indicating that combined SHP1/SHP2 deficiency did not interfere with proliferation. By contrast, at three hours post-TCR stimulation, DKO CD4+ T cells exhibited three-fold greater Annexin V positivity than controls (27.5% and 8.9%, respectively; Fig. 4F). No other genotype showed significant elevations in Annexin V binding on CD4+ T cells at this time point. Notably, the percentage of apoptotic CD8+ T cells was lower than in CD4+ T cells and was similar in all genotypes (Fig. 4G).

Excessive or chronic TCR stimulation can result in activation-induced cell death, an apoptotic process that contributes to peripheral tolerance (37, 38). Activation-induced cell death is mediated by FAS ligand (FASL) binding to FAS on T cells (39, 40). To test whether DKO CD4+ T cells were dying by FASL/FAS-mediated activation-induced cell death, we stimulated T cells with or without FASL-blocking antibody. After 4 days, FASL blockade increased control CD4+ T cell percentages from 24.6% to 33.8% (1.4 -fold), reflecting a rescue of normal activation-induced cell death in these cells (Fig. 4H). By contrast, FASL blockade increased DKO CD4+ T cell percentages from 5.6% to 21.9% (3.9-fold increase), accounting for 86% of the excess DKO CD4+ T cell death. Collectively, these results indicate that lack of both SHPs impairs CD4+ T cell expansion mainly by increasing FAS-mediated activation-induced cell death.

### Adoptive Transfer of Antigen-Specific CD4+ T Cells Restores Antitumor Responses in DKO Mice

To test whether the defective antitumor response in DKO mice resulted from loss of functional CD4+ T cells, we generated OVA-expressing MC38 cells (MC38-OVA) and injected mice subcutaneously on their right flanks with MC38- OVA and on their left flanks with parental MC38 cells. On the same day, we adoptively transferred control 2D2 CD4+ T cells (expressing a transgenic TCR specific for the unrelated MOG peptide) or wild type OT-II CD4+ T cells (expressing a transgenic TCR specific for OVA_323-339_ peptide) into control and DKO mice (Fig. 5A). As expected, parental MC38 tumors grew significantly larger in DKO mice than in control mice, regardless of 2D2 or OT-II CD4+ T cell treatment (Fig. 5B). Adoptive transfer of OT-II CD4+ T cells did not enhance antitumor responses against MC38-OVA tumors in control mice (Fig. 5C), arguing that tumor-infiltrating CD4+ cells are not limiting in these animals. By contrast, transfer of OT-II CD4+ T cells, but not 2D2 CD4+ T cells, restored the dysfunctional antitumor response in DKO mice (Fig. 5D). Thus, lack of competent CD4+ T cells can explain the dysfunctional antitumor response in DKO mice.

**Figure 5.**
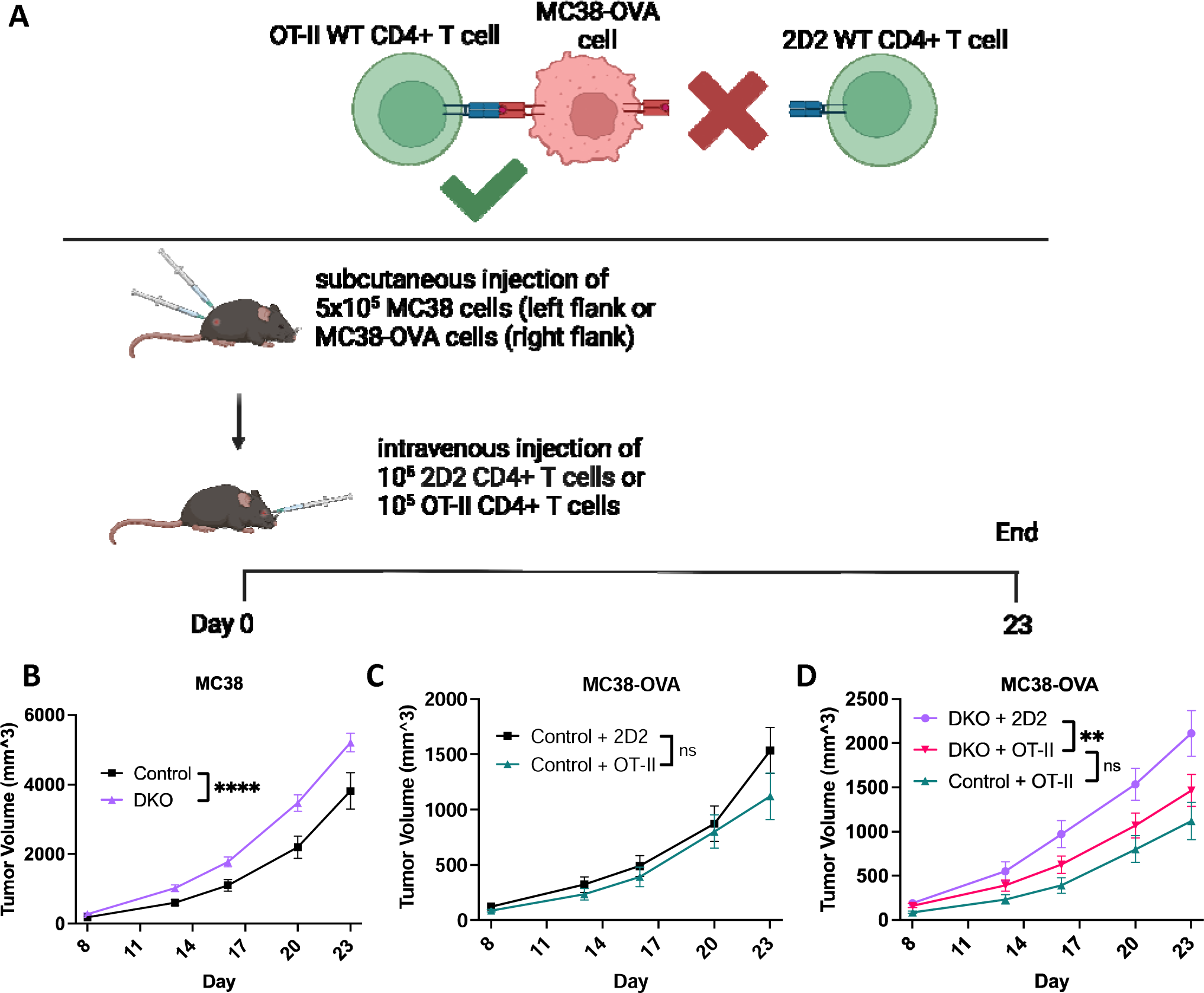
CD4+ T cell deficiency is responsible for dysfunctional DKO antitumor responses. (A) Diagram depicting experimental setup. (B-D) MC38 and MC38-OVA tumor growth in mice following adoptive transfer of 2D2 or OT-II CD4+ T cells. (B) MC38 tumors in WT (black, n=21) and DKO (red, n=32) mice, (C) MC38-OVA tumors in WT mice following adoptive transfer if 2D2 (black, n=10) or OT-II (blue, n=11) CD4+ T cells, (D) MC38-OVA tumors in DKO mice following adoptive transfer of 2D2 (green; n=16) or OT-II CD4+ T cells (red; n=16). All comparisons are combined from 2 experiments and analyzed by two-way ANOVA. Mean ± standard error are indicated. Not significant (ns), *p<0.05, **p<0.01, ***p<0.001, ****p<0.0001.

### DKO Mice Also Have Abnormal Responses to Acute Viral Infection

To determine whether DKO mice had impaired immune responses in a context other than tumors, we infected mice with Vaccinia-OVA (VACV-OVA) through scarification of their ears. Multiple reports indicate that CD4+ T cells are important for expansion of VACV-specific CD8+ T cells and/or elimination of virus, and previous work found that CD4+ T cell depletion impairs viral clearance at day 15 (41–43). We found that DKO mice had significantly increased ear thickness over the duration of the experiment (Fig. S5A), reflecting impaired resolution of the immune response. At day 15, 3/4 control mice had cleared infection, while the fourth only yielded 30 plaque forming units (PFU; Fig. S5B). By contrast, 2/4 DKO mice failed to clear the infection with 2,000 and 20,000 PFU. Moreover, total T cells were decreased in the ears of DKO mice relative to control mice, with CD4+ T cells being significantly lower and CD8+ T cells trending downward (Fig. S5C-E). Unfortunately, I-Ab OVA_232-339_ tetramers, used to identify OVA-specific CD4+ T cells, showed poor staining. However, there were fewer antigen-specific CD8+ T cells in the ear parenchyma of DKO mice compared with control mice (as measured by SIINFEKL H2-Kb tetramers; Fig. S6F). Collectively, these data are consistent with a defective CD4+ T cell response, impairing antigen-specific CD8+ T cell proliferation and clearance of virus.

## DISCUSSION

SHP1 and SHP2 have substantial structural and sequence similarity and share several binding proteins, suggesting that they could have some redundant functions. In T lymphocytes, however, the effects of deletion of each SHP are distinct: SHP1 KO mice have increased effector/central memory CD4+ and CD8+ populations and Th2 skewing, whereas SHP2 KO mice reportedly have expanded CD8+ T cells. To identify potentially redundant functions, we generated mice with combined SHP1/SHP2 deficiency. Our results reveal that the absence of both SHPs in T cells leads to increased peripheral memory T cell populations/Th2 skewing. Importantly, only SHP1/SHP2 double deficiency results in enhanced CD4+ T cell susceptibility to activation-induced cell death, and an impaired antitumor immune response.

Prior work has assessed the effects of individual deficiency of SHP1 and SHP2 on thymocyte development. Martinez et al. found that TCR signal strength was increased in SHP1 KO (*Cd4-Cre Ptpn6^/fl^*) thymocytes, leading to fewer CD4+ and CD8+ SP thymocytes compared with controls (6). Consistent with these findings, we observed increased markers of TCR signal strength and decreased numbers of DP and SP SHP1 KO thymocytes relative to control mice (Fig. S1). Notably, in an earlier study of T cell-specific SHP1 KO (*Cd4-Cre Ptpn6^/fl^*) mice performed at another institution, we failed to see any difference in the thymic compartment. The reason for this discrepancy is unclear; conceivably, differences in animal facilities play a role. One earlier report argued that SHP2 affects thymocyte development by limiting TCR signaling (44). However, these results might have been confounded by the use of a proximal LCK-Cre to drive *Ptpn11* deletion; notably, this driver itself causes a similar thymic phenotype (45). In agreement with work by Rota et al., who also analyzed *Cd4-Cre Ptpn11^/fl^* mice (14, 35), we observed no gross differences in thymocyte development in SHP2 KO mice. We also found that compound SHP1/SHP2 deficiency is similar to SHP1 deficiency except for a small decrease in DP thymocyte number in DKO mice. Our results contrast with those of Ventura et al., who recently reported that DKO mice (*Cd4-Cre Ptpn6^fl/fl^ Ptpn11^fl/fl^*) and *Ptpn6^fl/fl^ Ptpn11^fl/fl^* controls have similar numbers of DP thymocytes, although they also observed lower numbers of SP CD4+ and CD8+ thymocytes (35).

We reported previously (10) and confirm herein (Fig. 1) that SHP1 restrict differentiation of peripheral naïve CD4+ and CD8+ T cells to effector and central memory cells and inhibits polarization of CD4+ T cells toward the Th2 phenotype. In our earlier study, we attributed these effects to enhanced IL-4R signaling, as IL-4 deficiency or blockade reversed the phenotype. Notably, the phosphorylated ITIMs of IL-4Rl1l bind both SHPs, suggesting that SHP2 might be able to at least partially substitute for SHP1 in this pathway (46). Indeed, while we found that SHP2 deficiency alone may not affect naïve T cell differentiation, combined SHP1/SHP2 deficiency drives even more effector/memory cell differentiation relative to SHP1 deficiency alone (Fig. 1). We also observed that SHP1 KO mice have an increased CD8+ VM compartment, which is further enlarged in DKO mice. Consistent with this finding, IL-4R-deficient mice have reduced populations of CD8+ VM T cells (47). Notably, transgenic expression of the catalytically inactive mutant SHP2-C459S (purported to act as a dominant negative mutant) specifically in T cells resulted in increased proportions of CD44^hi^ T cells and enhanced expression of IL-4, IL-5, and IL-10 (48). Conceivably, this putative “dominant negative SHP2” mutant, in addition to competing with endogenous SHP2, also blocks SHP1 binding to IL4Rα, thereby mimicking the DKO phenotype.

We also determined the effects of SHP1, SHP2, and combined SHP1/SHP2 deficiency on antitumor responses using the well-characterized subcutaneous MC38 model (Fig. 2). Because PD-1, among other inhibitory receptors that bind the SHPs, is known to impair antitumor T cell responses, we asked whether PD-1 blockade could enhance antitumor T cell responses in these mice. Deletion of an essential mediator of PD-1 signaling should phenocopy PD-1 deletion or blockade, yielding smaller tumors regardless of treatment. Rota et al. reported that SHP2 KO mice respond to PD-1 blockade, indicating that T cell expression of SHP2 was not required for the anti-tumor effects of PD-1 antibodies (14). Not only did we confirm that T cell-SHP2 is dispensable for the effects of PD-1 blockade, but we observed that SHP2 deficiency in T cells actually enhances these effects via an as yet unclear mechanism (Fig. 2). Conceivably, SHP2 negatively regulates another pathway(s), whose effects are masked in the presence of intact PD-1 signaling. Indeed, while tumor growth in isotype-treated SHP2 KO mice and control mice was similar, 2/23 SHP2 KO mice qualified as responders despite having received no anti-PD-1 treatment (Fig. 2). This finding suggests SHP2 may indeed mediate signaling by a parallel inhibitory pathway(s). Notably, Rota et al. did not compare PD-1 treatment efficacy between SHP2 KO mice and CD4-Cre-expressing controls (14). Regardless of the precise underlying mechanism, our results argue that combining SHP2 inhibitors with PD-1 blockade might have therapeutic benefit and raise the question of whether SHP2 deficiency in T cells might sensitize tumors unresponsive to PD-1 blockade.

In agreement with a recent study, we found that SHP1 expression in T cells also is dispensable for the response to PD-1 blockade (35). However, while Ventura et al. found that PD-1 blockade was less effective at impeding tumor growth in SHP1 KO mice than in controls, we observed no difference in PD-1 response in our SHP1-deficient animals (Fig. 2) This discrepancy likely stems from their use of floxed littermate controls instead of the CD4-Cre-expressing controls used in our study. Cre has deleterious effects in multiple cell types, and we found that CD4-Cre mice have a blunted response to PD-1 blockade (Fig. S2). Like Ventura et al. we observed that DKO mice fail to respond to PD-1 blockade and have larger tumors than controls, indicating that DKO mice do not phenocopy PD-1 blockade or deficiency. These results also indicated that the antitumor T cell response was dysfunctional, making it impossible to determine whether PD-1 signaling in T cells requires SHP1 and/or SHP2. While our study was in progress, Strauss et al. reported that PD-1 inhibits antitumor immune responses in the MC38 model primarily by acting in myeloid cells, with only a smaller contribution by T cells (26). Thus, it remains to be determined whether either SHP1 or SHP2 is required for PD-1 function in T cells.

Tumors in DKO mice had decreased frequencies of CD4+ and CD8+ T cells as well as impaired activation of dendritic cells (Fig. 3). These findings comport with a model in which loss of CD4+ T cells leads to decreased licensing of dendritic cells and results in suboptimal CD8+ T cell activation. Notably, tumors from DKO mice contained less than a third of the number of dendritic cells observed in controls (Fig. 3). They also had increased percentages of TAMs, and these TAMs expressed lower levels of PD-L1 than TAMs in control mice. Previous studies showed that GM-CSF produced by T cells promotes upregulation of PD-L1 on TAMs and recruitment of dendritic cells (49–53). CD4+ T cells also promote recruitment of CD8+ T cells (Bos & Sherman, 2010), which could help explain the lower frequencies of CD8+ T cells in tumors from DKO mice

Combined deficiency of SHP1 and SHP2 increased activation-induced cell death in CD4+ T cells *in vitro* (Fig. 4), potentially explaining the decrease in CD4 T cells in tumors from DKO mice. Importantly, the increased activation-induced cell death of CD4+ T cells from DKO mice cannot be explained by the increased percentage of effector T cells in these mice: SHP1 KO mice have similar proportions of splenic effector CD4+ T cells but did not show increased rates of apoptosis upon stimulation. High and repeated antigenic stimulation *in vivo*, such as would occur in MC38 tumor-bearing mice, results in deletion of reactive clones (40, 54). *Fas^lpr/lpr^* (Lpr) mice, which express extremely low levels of FAS, have T cells incapable of activation-induced cell death, leading to autoimmunity (39, 40, 55). FAS blockade largely rescues CD4+ T cell death *in vitro*, as expected if these cells are undergoing activation-induced cell death (Fig. 4). Notably, THEMIS forms a complex with both SHPs, and THEMIS knockdown in primary human peripheral CD4+ T cells results in significantly enhanced activation-induced cell death upon TCR stimulation (56, 57). Conceivably, SHP1 and SHP2 both act via THEMIS to inhibit TCR signaling and prevent activation-induced cell death. It will be interesting to determine whether THEMIS knockout mice also have enlarged tumors and impaired anti-PD-1 response compared with controls. Most importantly, we found that adoptively transferring normal CD4+ T cells to DKO mice restored their antitumor response (Fig. 5), consistent with the idea that CD4+ T cell deficiency is a major reason for impaired tumor control.

In contrast to the effects of combined SHP1/SHP2 deficiency on CD4+ T cells, CD8+ T cells from DKO mice showed no increase in activation-induced cell death. Control CD8+ T cells also showed lower levels of activation-induced cell death than control CD4+ T cells, consistent with previous results (58). By contrast, Ventura et al. reported decreased CD8+ T cell expansion in their DKO mice. Notably, the two studies monitored T cell expansion over different durations, used different Cre driver strains (CD4-Cre versus GZMB-Cre), and employed different controls (Cre-expressing controls here versus floxed controls in their report) (35), any or all of which could explain these discrepancies. Although our data suggest a more dominant role of SHP1/2 double deficiency on CD4 T cells, it is possible that it may also have direct effects on CD8 T cells and play important function in tumor immunity, especially in different tumor models beyond MC38.

In summary, we conclude that SHP1 and SHP2 serve an indispensable role in CD4+ T cell function. T cell deficiency of both PTPs in mice results in severely impaired T cell-mediated antitumor responses, including a failure to respond to PD-1 blockade. Tumors in DKO mice have a dysregulated immune microenvironment, characterized by changes consistent with loss of CD4+ T cell help. CD4+ T cells, but not CD8+ T cells, from DKO mice undergo increased FAS-dependent apoptosis in response to TCR stimulation, and loss of normal CD4+ T cell function can account for the impaired antitumor T cell responses in DKO mice. DKO mice similarly show dysregulated immune responses when challenged with VACV-OVA. (Fig. S5) Collectively, our findings provide insight into how SHP1/SHP2 regulate CD4+ T cell survival upon activation and suggest new therapeutic opportunities for patients with cancer.

## MATERIALS AND METHODS

### Cell lines

MC38 cells were maintained at 37°C in 5% CO_2_ in DMEM containing 10% FBS and penicillin plus streptomycin (Thermo Fisher Scientific; 15140163). To generate MC38-αCD3 cells, we cloned the scFv sequence of murine-reactive αCD3 clone 145-2C11 into the CD5L-OKT3-scFv-CD14 lentiviral plasmid generated by Leitner *et al.*, replacing the OKT3 region

(59). Lentiviral particles were generated by transfecting HEK 293T cells with lentiplasmid, psPAX2, and pMD2.G plasmids. MC38 cells were then transduced with CD5L-145-2C11-scFv-CD14 lentivirus and purified by FACS for human CD14. MC38-OVA cells were generated by transducing MC38 cells with pLVX-puro-cOVA (Addgene plasmid #135073) followed by selection in puromycin (4 µg/mL).

### Mice

All animal studies were approved by the Institutional Animal Care and Use Committee at New York University Grossman School of Medicine. *Ptpn6^fl/fl^* and *Ptpn11^fl/fl^* mice were described previously and were maintained on C57Bl/6 background

(60). CD4-Cre-expressing mice were provided by Christopher B. Willson (30).

### Immunoblotting

Splenic T cells isolated by negative selection were stained with anti-CD45 FITC and anti-CD3 BV421 (BioLegend; 100336) and CD45+ CD3+ cells were recovered by FACS and centrifugation. Pellets were lysed in 1% NP-40 buffer with 0.1% SDS supplemented with phosphatase inhibitors (10 nM NaF, 1 mM Na_3_VO_4_, 10 mM b-glycerophosphate, 10 mM sodium pyrophosphate) and proteases (40 µg/mL PMSF, 2 µg/mL antipain, 2 µg/mL pepstatin A, 20 µg/mL leupeptin, and 20 µg/mL aprotinin), and clarified by centrifugation at 20,000 rcf for 10 minutes at 4 °C in a microfuge. Protein was quantified by Coomassie Blue assay (Pierce™ Bradford Protein Assay Kit; 23200). Lysates were boiled in SDS-PAGE sample buffer, resolved on 10% SDS-PAGE gels, and transferred onto PVDF membranes in transfer buffer (glycine 14.4 g/L, Tris 3.03 g/L, 15% methanol; pH 8.6). Membranes were blocked with 2% milk, incubated with primary antibodies [anti-SHP1 (Santa Cruz; sc-287; 1:000), anti-PTPN11 (Santa Cruz; sc-7384; 1:1000), and anti-ERK2 (Santa Cruz; sc-1647; 1:1000)] at 4 °C overnight in 5% BSA in TBST, washed in TBST, and incubated with secondary antibodies [goat anti-rabbit 680LT (LI-COR; 926-68021; 1:20,000) and goat anti-mouse 800CW (LI-COR; 926-32210; 1:20,000)] for 1 hour at room temperature. Membranes were washed in TBST and dried before imaging on the LI-COR Odyssey CLx. Signals were quantified using Image Studio from LI-COR.

### *Ex vivo* T cell stimulations and killing assays

For cell expansion assays, splenic T cells were isolated by negative selection (STEMCELL Technologies; 19851) and added to 96-, 24, or 6-well plates [pre-coated overnight at 4 °C with goat anti-hamster IgG (MP Biomedicals; 856984)] in T cell stimulation medium containing anti-CD3 (1 µg/mL; BioLegend; 100238) and anti-CD28 (1 µg/mL; BioLegend; 102116) at 10^6^ cells/mL (1 mL for 24-well plates and 3 mL for 6-well plates) or 2*10^6^ cells/mL (200 µL for 96-well plates). After incubation for 48 hours, cells were resuspended in fresh T cell medium [RPMI with 10% FBS, penicillin plus streptomycin (Thermo Fisher Scientific; 15140163; 100 U/mL and 100 mg/mL), 20 mM HEPES, GlutaMAX (Thermo Fisher Scientific), fresh 2-mercaptoethanol (55 µM) and recombinant human IL-2 (20 ng/mL; STEMCELL Technologies; 78036)] and transferred to a new plate to discontinue stimulation. T cells were counted and resuspended in T cell medium daily at 1-3x10^6^ cells/mL. T T cells were stained at 3 hours post-stimulation with Annexin V (BD; 559763) according to manufacturer’s instructions and anti-CD4 BUV737 and aCD8 BV785 (BioLegend; 100750). T cells were also stained with anti-CD4 BUV737 and anti-CD8 BV785 on days 4 and 5. For killing assays, MC38 or MC38-αCD3 cells were added at 5x10^4^ cells/well to 96-well RTCA E-plates (Agilent; 300600910). On day 7 post-T cell stimulation, T cells were added at various E:T ratios to the wells in. Killing was measured by adherent cell impedance every 15 minutes. Impedance at each time point was normalized to the impedance value at time of T cell addition.

### Lymphoid tissue characterization

Thymuses/spleens/bilateral inguinal LNs were collected from 4-week-old (thymus) or 9-11 week-old (spleen LNs) age-and sex-matched mice, weighed (spleen/thymus), mechanically dissociated, and washed by passage through a 70 µm strainer, treated with 180 µL DNase I (STEMCELL Technologies; 07900; 1 mg/mL stock) in 3 mL FACS buffer [2% fetal bovine serum in phosphate-buffered saline (PBS)] with 5 mM MgCl_2_ for 15 minutes (thymus only), washed, and treated with ACK lysis buffer for 3 minutes. After washing again, viable cells were counted using trypan blue exclusion. Cells (10^6^) were treated with mouse FcR blocking reagent (Miltenyi; 130-092-575) and stained with appropriate panel (see Supplementary Tables 1 and 2) for 30 minutes in the dark at 4 °C. Samples were washed in PBS and stained with LIVE/Dead Fixable Blue (Thermo Fisher Scientific; L23105) for 15 minutes in the dark at 4 °C, washed, fixed in 2% paraformaldehyde for 20 minutes in the dark at 4 °C, and washed again before acquiring data on an LSR II UV flow cytometer. CD5 and CD69 expression was assessed on samples gated on cells, single cells, live cells, B220-cells, and CD4+CD8- or CD4-CD8+ cells. CD5 and CD69 gates were set based on fluorescence minus one controls (FMOs). CD44, CD62L, and CD49d expression was assessed on samples gated on cells, single cells, live cells, CD45+ cells, and CD4+CD8-or CD4-CD8+ cells. The CD49d gate was set based on FMOs.

### Tumor assays

Mice (8-12-week-old, both male and female, sex-matched) were injected subcutaneously with 5x10^5^ MC38 or MC38-OVA cells in 100 µL PBS on day 0. Where indicated, mice were treated with 200 µg anti-PD-1 (Bio X Cell; BE0146; clone RMP1-14) or isotype control antibody (Bio X Cell; BE0089; clone 2A3) on days 5, 8, 11, and 14. Tumors were measured by using calipers, and volume was calculated as: V = 0.5(L x W^2^) where V=volume, L= length of longest dimension, and W = length of perpendicular dimension. Similar to prior studies (61–63), we sorted mice into responders and non-responders. We defined responders as mice whose tumors had a less than 30% increase in volume between day 8 (the first day of measurement) and day 23 (the last day of measurement). Non-responders were defined as mice whose tumors exhibited a ≥30% increase in volume. This cutoff represents a stringent threshold, as even a 1 mm fluctuation in each tumor dimension due to caliper measurement error could result in apparent tumor volume changes exceeding 50% at the average control mouse tumor size on day 8 (116 mm³). Mice humanely sacrificed for tumor ulceration or poor health before day 23 were excluded from statistical analysis (by chi-square test). Groups of mice were only compared if from the same experiment (e.g., isotype vs. l1lPD-1, isotype vs. isotype, l1lPD-1 vs. l1lPD-1). For adoptive transfer of wild type 2D2 or OT-II CD4+ T cells, donor splenic T cells were isolated by negative selection (STEMCELL Technologies; 19852), and 10^5^ cells were injected retro-orbitally in PBS into sex-matched recipients after tumor injection on day 0.

For characterization of the immune microenvironment, mice were anesthetized with ketamine and perfused with 30 mL ice-cold PBS containing 2-5 mM EDTA to flush out leukocytes remaining in the vasculature. Tumors were weighed, and a portion of tumor was placed in 2 mL digestion mix [9 parts DMEM, 1 part 10x collagenase/hyaluronidase (STEMCELL Technologies; 07912), 0.1 parts DNase I], cut into 1-2 mm pieces, and incubated for 30 minutes at 37 °C and 5% CO_2_ with intermittent gentle swirling. Digested tumors were pressed through 70 µm strainers, washed, and treated with ACK lysis buffer for 1 minute before washing again. Cells were counted, and cells (1.5x10^6^) were treated with mouse FcR blocking reagent for 10 minutes at 4 °C and stained in the dark at room temperature for 20 minutes with the appropriate panels (see Supplementary Tables 1 and 2). Aliquots were also treated with 81 nM PMA, 1.39 µM ionomycin, and 5 µg/mL Brefeldin A and incubated at 37 °C, 5% CO_2_ for 4 hours before staining with P4 (see Supplementary Tables 1 and 2). Samples were washed in PBS, stained with LIVE/Dead Fixable Blue (Thermo Fisher Scientific; L23105) for 15 minutes in the dark at 4 °C, washed again, fixed overnight (eBioscience FoxP3 transcription factor staining buffer set; Thermo Fisher scientific; 00-5523-00), and permeabilized. Samples were then incubated with FcR blocking reagent, rat serum, and mouse serum for 10 minutes before staining with panel antibodies targeting intracellular antigens for 30 minutes at 4°C. For P1, CD69, LAG3, TIM3, PD-1, OX40, and Granzyme B expression was assessed on samples gated on cells, single cells, live cells, CD45+ cells, CD3+ cells, and CD4+CD8-or CD4-CD8+ cells. Gates for CD69, Granzyme B, LAG3, OX40, TIM3, and Reps1 dextramer were set based on FMOs. For P2, samples were gated on cells, single cells, live cells, CD45+ cells, and CD19+ (B cells) or CD19-cells. From the CD19-population, cells were gated on MHC-II expression. MHC-II+ cells were gated as F4/80+CD11c+ (TAMs) or F4/80-CD11c-(dendritic cells). MHC-II-cells were gated on CD11b expression. CD11b+ cells were gated as Ly6C+Ly6G-(M-MDSCs) or Ly6C+Ly6G+ (G-MDSCs). CD49b+ (NK cells) were gated from the CD11b-population. FMO was used for PD-L1 gating. For P3, CD44, CD69, CTLA4, FOXP3, T-BET, and RORγt populations were assessed by gating on cells, single cells, live cells, CD45+ cells, CD4+CD8- or CD4-CD8+ cells, and CD3+ cells. FMOs were used for gating CD69, CTLA4, RORγt, T-BET, and GATA3. For P4, IFNγ, TNFα, and IL-2 expression were assessed by gating on cells, single cells, live cells, CD45+ cells, CD3+CD8+ cells. FMOs were used to gate on IFNγ, TNFα, and IL-2.

### Vaccinia virus infection

VACV-OVA was propagated in BSC-40 cells. On day 0, mice were infected by adding 10 µL of 5x10^6^ PFU of VACV-OVA to the ventral side of each pinna and subsequently scarifying with a 29-gauge needle. Ear thickness was measured by caliper on days 0, 7, 9, 11, and 15. Five minutes before sacrifice, mice were retro-orbitally injected with aCD45 BUV395 (2 µg) to label leukocytes still in the vasculature (for analytical exclusion). On day 15, mice were euthanized, and one ear was taken for plaque assay. The other ear was collected for flow cytometry. Dorsal and ventral sides of the ear were separated and incubated at 37 °C for 45 minutes in HBSS (Hyclone) containing CaCl_2_, MgCl_2_, 125 U/mL collagenase D (Invitrogen), and 60 U/mL Dnase-I (Sigma-Aldrich). Ears were ground on a scored plastic dish and poured through a 70 µm strainer. Cells (1.5x10^6^) were stained with H-2Kb SIINFEKL (OVA_257-264_) tetramer BV421 (NIH tetramer core facility) for 45 minutes at room temperature. Samples were washed, FcR-blocked, and stained with anti-CD3 BV605, anti-CD4 BUV737, and anti-CD8a BV785. Samples were then stained with LIVE/Dead Fixable Blue for 15 minutes in the dark at 4°C, washed, fixed in 2% paraformaldehyde for 20 minutes in the dark at 4 °C, and washed again before acquiring data on a BD FACSymphony A5 flow cytometer.

### Quantification and Statistical Analysis

Graphs were generated and statistical analyses were performed using GraphPad Prism v.10. Groups were compared by two-way unpaired t-tests, one-way ANOVA, two-way ANOVA, or chi-squared test as indicated. Area under the curve was calculated with RStudio. Means with error bars representing standard error of the mean are shown. Relevant *P* values are indicated in the figures.

### Illustrations

Illustrations were created with BioRender.

## ACKNOWLEDGEMENTS

We thank members of the Wang and Neel labs for helpful discussions. We also thank Dr. Stefan Feske (NYU School of Medicine) and Dr. Christopher B. Wilson (University of Washington) for sharing mice. This work was supported in part by NIH grants R01CA248896 (B.G.N.), R37CA273333–01 (J.W.), R01CA269898 (J.W.), R01AR080068 (A.W.L.), T32GM007308 (C.J.R.F), T32CA009161 (C.J.R.F), T32AI100853 (T.H.), and a young investigator award from the Melanoma Research Alliance (J.W.). Flow cytometry was supported by P30CA016087.

---

**Figure S1.**
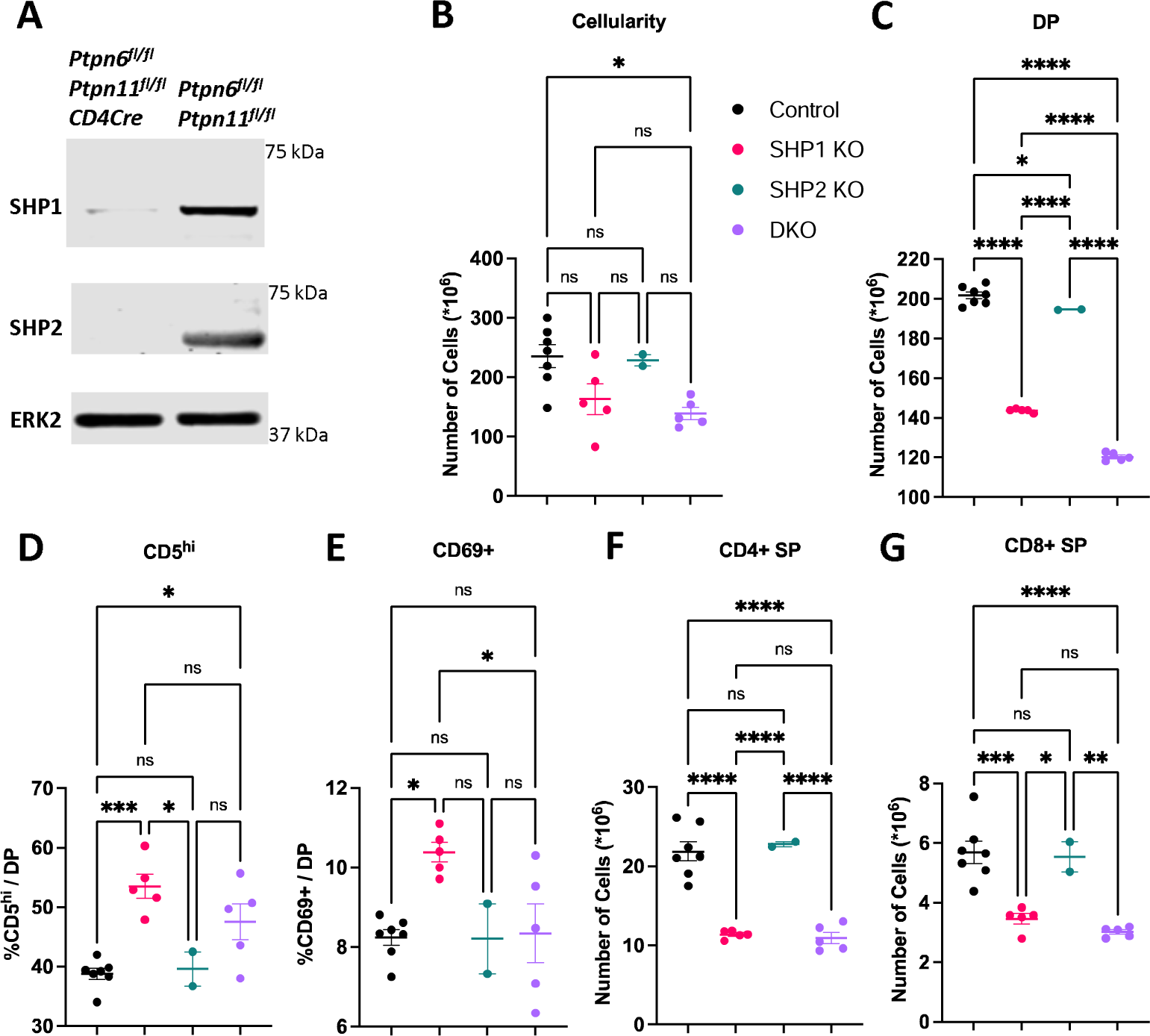
Thymocyte development is altered in SHP1 KO and DKO mice. (A) Immunoblot showing SHP1, SHP2, and ERK (loading control) levels in splenic T cells from the indicated mice. A single experiment is shown. (B) Thymic cellularity, (C) CD4+ CD8+ (DP) thymocyte number, (D-E) DP cells stained for CD5 (D) or CD69 (E), as indicated. (F-G) Number of CD4 (G) and CD8 (H) single positive (SP) thymocytes. Data are combined from two experiments. Each point represents value from one mouse. Means ± standard error are indicated. Significance was evaluated by one-way ANOVA with multiple comparisons, not significant (ns), *p<0.05, **p<0.01, ***p<0.001, ****p<0.0001.

**Figure S2.**
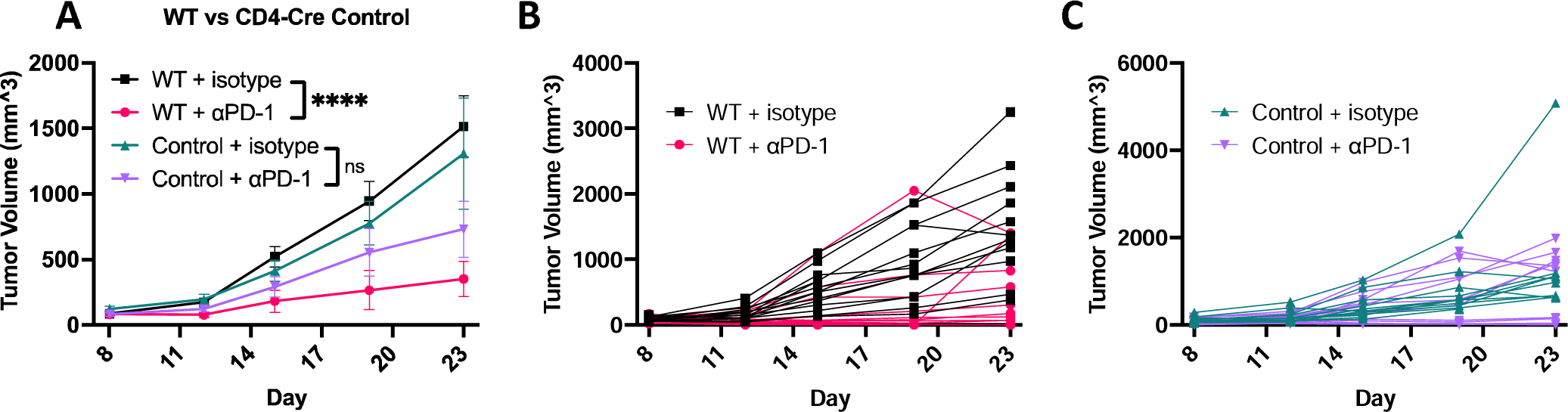
CD4-Cre impairs antitumor T cell responses. (A-C) MC38 tumor growth in *Ptpn6^+/+^ Ptpn11^+/+^* + isotype antibody (green; n=14), *Ptpn6^+/+^ Ptpn11^+/+^* + anti-PD-1 (bluem n=14), *Cd4-Cre Ptpn6^+/+^;Ptpn11^+/+^* + isotype antibody (black, n=12), and *Cd4-Cre Ptpn6^+/+^;Ptpn11^+/+^* + anti-PD-1 (red, n=15) mice. (B-C) individual tumor growth curves for (B) *Ptpn6^+/+^;Ptpn11^+/+^* mice and (C) *Cd4-Cre Ptpn6^+/+^;Ptpn11^+/+^* mice. Tumor growth was analyzed by two-way ANOVA. All comparisons are combined from 2 experiments. Means ± standard error are indicated. Not significant (ns), *p<0.05, **p<0.01, ***p<0.001, ****p<0.0001.

**Figure S3.**
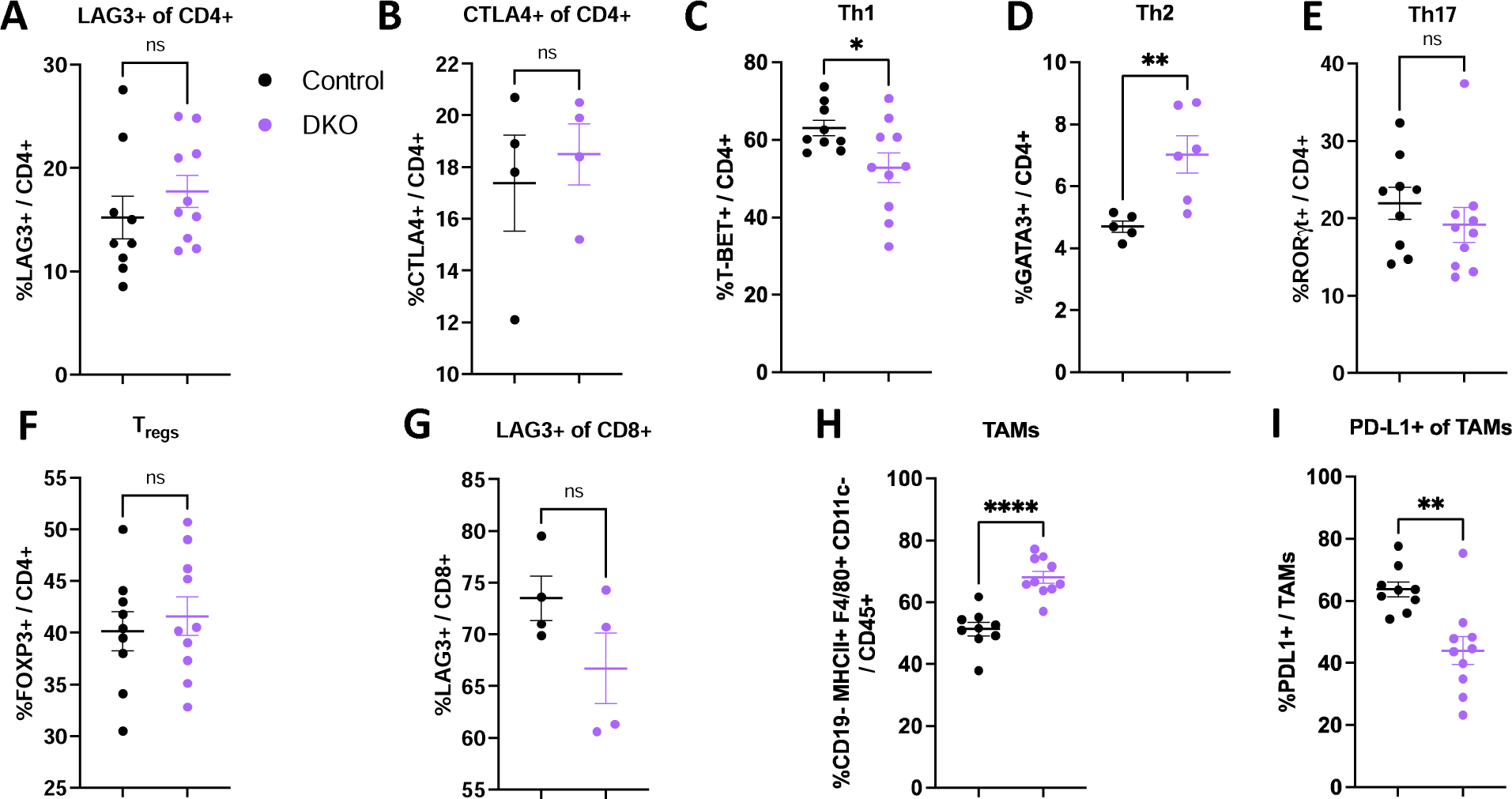
DKO mouse tumors have a dysregulated immune microenvironment (continued). (A-I) Flow cytometric analysis of isotype-treated WT and DKO mouse tumors. (A) LAG3+ CD4+ T cells, (B) CTLA4+ CD4+ T cells, (C) Th1 CD4+ T cells, (D) Th2 CD4+ T cells, (E) Th17 CD4+ T cells, (F) T_regs_, (G) LAG3+ CD8+ T cells, (H) tumor-associated macrophages, (I) PD-L1+ TAMs. Data are combined from two experiments. Each point represents value from one mouse. Means ± standard error are indicated. Significance was evaluated by two-tailed unpaired t-tests, not significant (ns), *p<0.05, **p<0.01, ***p<0.001, ****p<0.0001.

**Figure S4.**
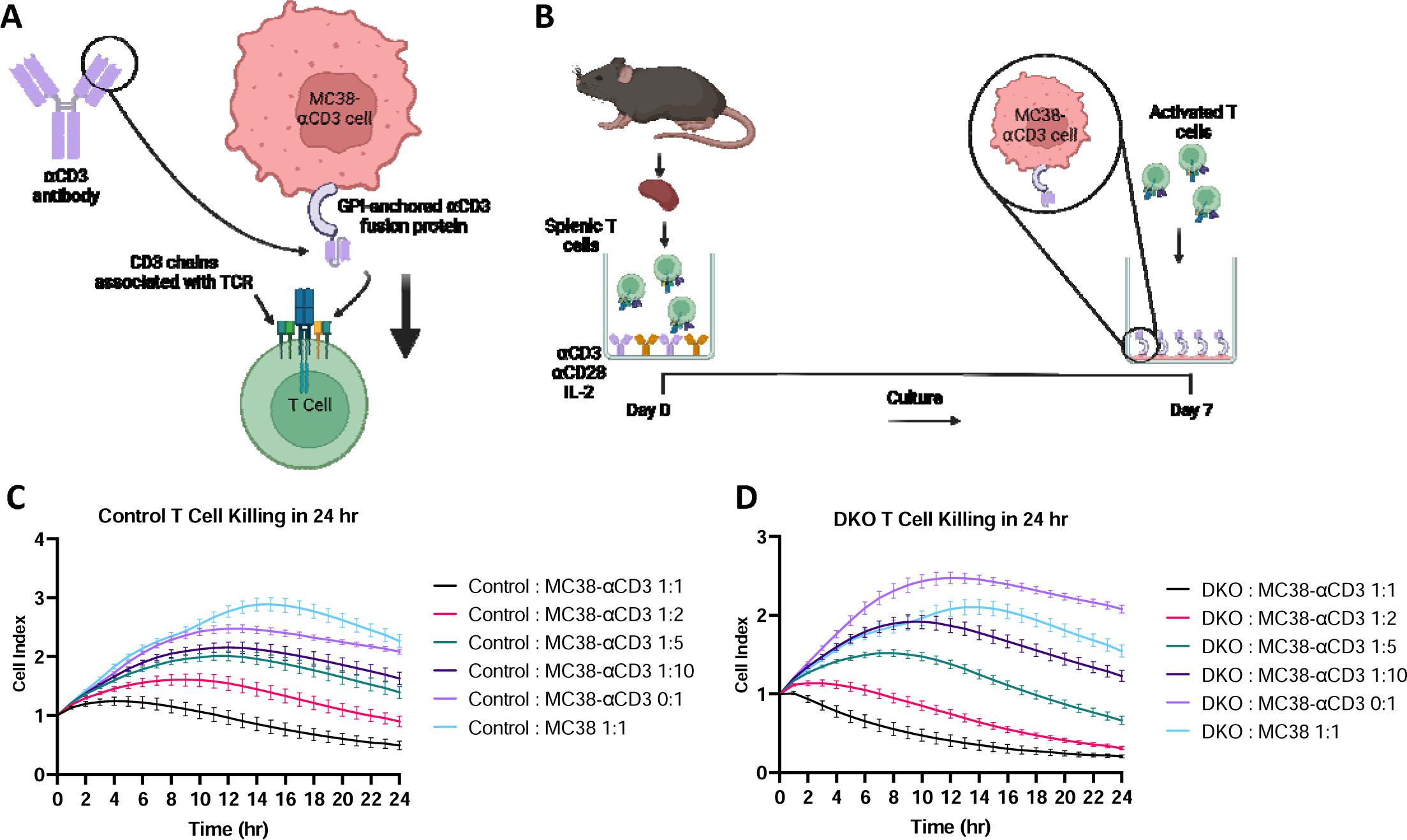
SHP1 and SHP2 are not required for T cell cytotoxicity. (A) Diagram depicting design of MC38-⍺CD3 cells. (B) Diagram depicting experimental setup. (C-D) Killing assays for co-cultures of (C) WT (n=4) or (D) DKO (n=4) T cells with MC38 or MC38-⍺CD3 cells at the indicated ratios. Area under the curve (AUC) was calculated for each biological replicate, and a two-tailed unpaired t-test was performed. Data are from one experiment. Means ± standard error are indicated. Not significant (ns), *p<0.05, **p<0.01, ***p<0.001, ****p<0.0001.

**Figure S5.**
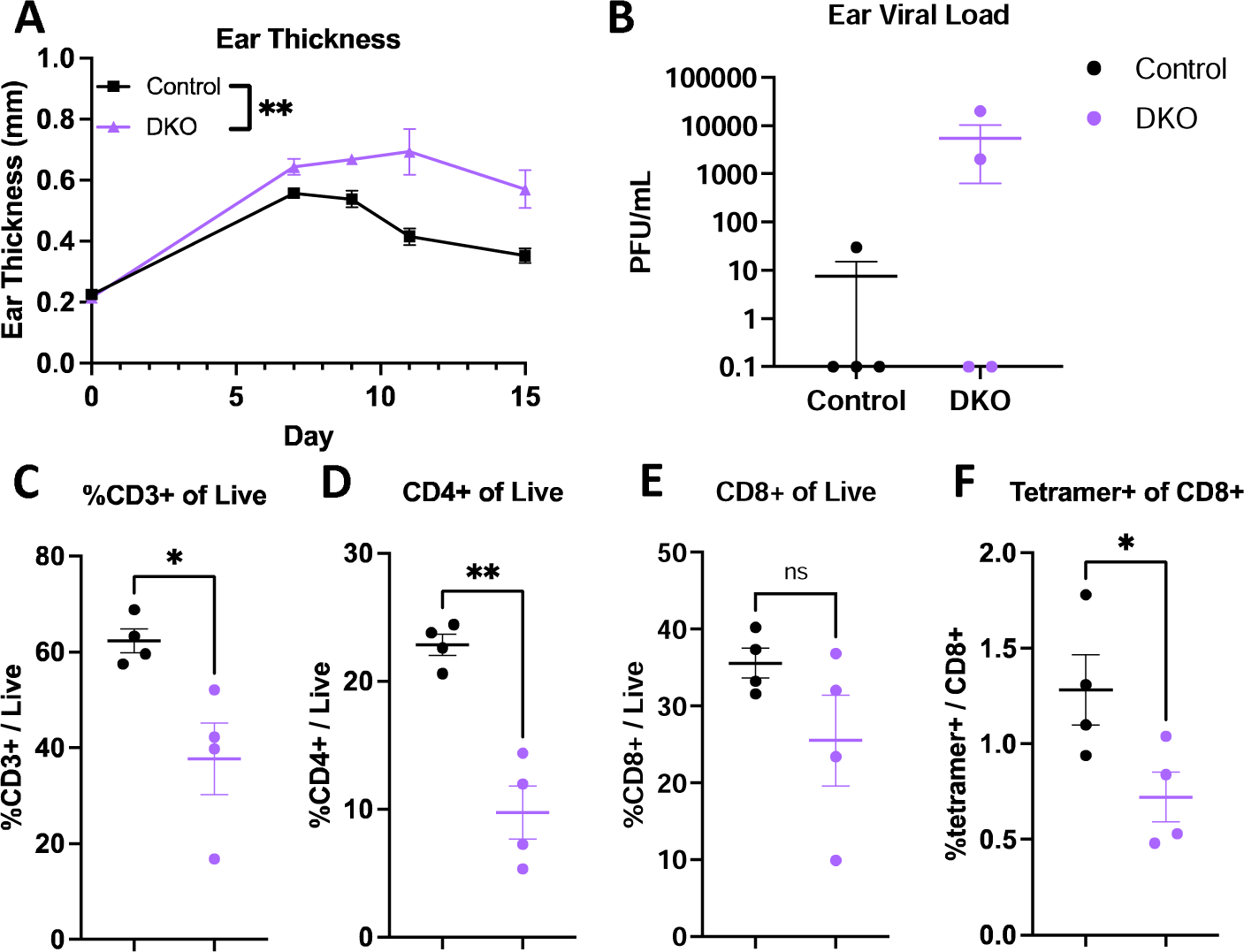
DKO mice have an impaired immune response to VACV-OVA. (A-F) Mice were infected with equal amounts of 1-5 x10^6^ plaque-forming units (PFU) of Vaccinia-OVA by scarification of the ear. (A) Ear thickness over time, analyzed by two-way ANOVA, (B) PFU from ears at day 15 post-infection, (C-F) Flow cytometric analysis of ear day 15 post-infection, (C) %CD3+ of live cells, (D) %CD4+ T cells of live cells, (E) %CD8+ T cells of live cells, (F) %SIINFEKL tetramer+ CD8+ T cells. Comparisons were analyzed by two-tailed unpaired t-tests. Data are from one experiment. Each point represents value from one mouse. Mean ± standard error are indicated. Significance was evaluated by one-way ANOVA with multiple comparisons, not significant (ns), *p<0.05, **p<0.01, ***p<0.001, ****p<0.0001.

**Supplementary Table 1:** Complete Flow Cytometry Antibody List.

| **Supplementary Table 1: Complete Flow Cytometry Antibody List** |  |
| --- | --- |
| **Antibody** | **Company and Catalog Number** |
| **anti-B220 APC/Fire 750** | BioLegend; 103259 |
| **anti-CD11b BV421** | BioLegend; 101236 |
| **anti-CD11c BUV737** | BD; 612796 |
| **anti-CD19 BV650** | BioLegend; 115541 |
| **anti-CD206 PerCP/Cy5.5** | BioLegend; 141716 |
| **anti-CD3 BV421** | BioLegend; 100228 |
| **anti-CD3 BV605** | BioLegend; 100237 |
| **anti-CD3 FITC** | BioLegend; 100204 |
| **anti-CD44 APC/Cy7** | BioLegend; 103028 |
| **anti-CD45 BUV395** | BD; 564279 |
| **anti-CD49b PE** | BioLegend; 108908 |
| **anti-CD49d PE** | BioLegend; 103608 |
| **anti-CD4 BUV737** | BD; 612844 |
| **anti-CD5 BB515** | BD; 565504 |
| **anti-CD62L PerCP/Cy5.5** | BioLegend; 104432 |
| **anti-CD69 PE/Cy7** | BioLegend; 104512 |
| **anti-CD69 PerCP/Cy5.5** | BioLegend; 104522 |
| **anti-CD8a BV570** | BioLegend; 100740 |
| **anti-CTLA4 APC** | BioLegend; 106310 |
| **anti-F4/80 APC/Cy7** | BioLegend; 123118 |
| **anti-FOXP3 PE** | Invitrogen; 12-5773-82 |
| **anti-GATA3 PE/Cy7** | BioLegend; 560405 |
| **anti-Granzyme B PE** | Invitrogen; 12-8899-41 |
| **anti-IFNγ BV650** | BioLegend; 505832 |
| **anti-IL-2 BV711** | BioLegend; 503837 |
| **anti-iNOS APC** | Invitrogen; 17-5920-82 |
| **anti-LAG3 BB515** | BD; 564672 |
| **anti-Ly6C BV570** | BioLegend; 128030 |
| **anti-Ly6G BV711** | BioLegend; 127643 |
| **anti-MHC-II PE/Cy7** | BioLegend; 107630 |
| **anti-OX40 PerCP/Cy5.5** | BioLegend; 119425 |
| **anti-PD-1 BV711** | BioLegend; 135231 |
| **anti-PD-L1 BV785** | BioLegend; 124331 |
| **anti-T-BET PE/Dazzle 594** | BioLegend; 644828 |
| **anti-TCF-1 BV421** | BD; 566692 |
| **anti-TIM3 BV785** | BioLegend; 119725 |
| **anti-TNFα BV785** | BioLegend; 506341 |
| **anti-RORγt BV650** | BD; 564722 |

**Supplementary Table 2:** Flow Cytometry Antibody Panels.

| **Panel** | **Extracellular Stains** | **Intracellular Stains** |
| --- | --- | --- |
| **Thymocyte** | anti-B220 APC/Fire 750, anti-CD4 BUV737, anti-CD8a BV570, anti-CD5 BB515, anti-CD69 PerCP/Cy5.5 |  |
| **Spleen/LN** | anti-CD45 BUV395, anti-CD3 BV421, anti-CD4 BUV737, anti-CD8a BV570, anti-CD44 APC/Cy7, anti-CD62L PerCP/Cy5.5, anti-CD49d PE |  |
| **P1** | anti-CD45 BUV395, anti-CD3 BV605, anti-CD4 BUV737, anti-CD8a BV570, anti-PD-1 BV711, anti-LAG3 BB515, anti-TIM3 BV785, anti-OX40 PerCP/Cy5.5, anti-CD69 PE/Cy7 | anti-TCF-1 BV421, anti-Granzyme B PE |
| **P2** | anti-CD45 BUV395, anti-CD19 BV650, anti-CD49b PE, anti-CD11b BV421, anti-CD11c BUV737, anti-Ly6C BV570, anti-Ly6G BV711, anti-F4/80 APC/Cy7, anti-MHC-II PE/Cy7, anti-PD-L1 BV785 | anti-CD206 PerCP/Cy5.5, anti-iNOS APC |
| **P3** | anti-CD45 BUV395, anti-CD3 BV605, anti-CD4 BUV737, anti-CD8a BV570, anti-CD44 APC/Cy7, anti-CD62L PerCP/Cy5.5, anti-CTLA4 APC | anti-FOXP3 PE, anti-T-BET PE/Dazzle 594, anti-GATA3 PE/Cy7, anti-RORγt BV650 |
| **P4** | anti-CD45 BUV395, anti-CD3 FITC, anti-CD8a BV570 | anti-IFNγ BV650, anti-TNFα BV785, anti-IL-2 BV711 |

